# Size matters, so does condition: the use of a body condition index reveals the costs and benefits of structural body size in an insect

**DOI:** 10.1101/774893

**Authors:** Caroline Zanchi, Yannick Moret, Mark A. F. Gillingham

## Abstract

1. Insects are core actors for the balance of many earth ecosystems, as well as an alternative source of food and feed with a low ecological footprint. A comprehensive understanding of their life history requires reliable tools. Body condition constitutes the amount of energy reserves available to a fitness trait after maintenance costs have been accounted for. Body condition is standardly estimated using Body Condition Indexes (BCIs) in vertebrates. In insects the relevance of BCIs is frequently questioned on the basis that they might not accurately reflect neither energy reserves nor fitness.. However, to date no study has tested whether the very concept of body condition is relevant in insects, i.e. whether BCIs accurately reflect the relative energy reserves allocated to fitness traits.
2. We propose that the relevance of using BCIs in insects depends on whether their structural size has a fitness cost. If on the contrary insects only benefit from a larger body size, a simple measurement of body size or mass will predict fitness, but not a BCI. We experimentally manipulated food availability at the larval and adult stage and used total fecundity of females as a fitness proxy of *Tenebrio molitor*, an important model in physiology, ecology and evolution, and one of the first insects to be considered as a source of food and feed.
3. Our results support three key assumptions of the relevance of BCIs in insects: (i) a valid BCI correlated with energy reserves corrected for a given size (i.e. relative energy reserves) and not with absolute measures of energy reserves; (ii) both structural size and body condition positively predict different components of fitness; and, (iii) the effect of body condition was dependent on resource availability, whereby its effect was only apparent and large when food was unrestricted at the larval stage and restricted at the adult stage.
4. Overall we demonstrate the relevance of using BCIs in insects. Their use should be generalized to improve fitness readouts in evolution, ecological and physiological studies, as well as improve their husbandry for commercial purposes.

## Introduction

Insects make up for the majority of species on the planet and might represent the highest biomass of terrestrial animals (Bar-On et al., 2018). The ease of their maintenance and manipulation coupled with their diverse life history has resulted in several insect species becoming prominent models in physiology (Scully & Bidochka, 2006; Schneider & Chambers, 2008; Keil & Steinbrecht, 2010; Adamski et al., 2019) and in ecology and evolution (Funk et al., 2002; Boggs, 2009; Bilton et al., 2019). Moreover, there is growing interest of cultivating insects as a source of food and feed (Kim et al., 2019; van Huis, 2021). These fields often require the measurement, sometimes repeated, of fitness traits during an insect’s lifespan, which is likely to be dependent on the amount of energy reserves present in individuals. For instance, the tracking of temporal variations in the nutritional state of experimental individuals could also allow the optimization of mass-rearing conditions in an industrial context (Gligorescu et al., 2020; Adámková et al., 2020). The amount of reserves present in an animal can be estimated by the direct -and invasive-quantification of several body components from the organism, such as fat, glycogen and protein content, which prevents the assessment of their predictive power on fitness and to follow their evolution. To circumvent this, **Body Condition Indexes (BCIs)** can be used.

BCIs are expected to reflect **body condition**, which is classically defined as the amount of reserves present in the body of an animal after maintenance costs have been accounted for (Rowe & Houle, 1996; Barnett et al., 2015). It is also sometimes referred to as the “general health state”, “physiological state”, or “quality” of an animal (Jakob et al., 1996; Wilson & Nussey, 2010). The relationship between the amount of reserves and the “quality” of an animal is likely to be more complicated than stated in most studies (Raubenheimer & Simpson, 2012; Barnett et al., 2015), which is why Hill (2011) defined body condition as being “the relative capacity to maintain optimal functionality of essential cellular processes” in an animal. In numerous cases, this capacity relies strongly on the amount of energy reserves an individual has at its disposal (Schulte-Hostedde et al. 2005; Nandy et al., 2012), which has been found to predict several fitness traits in animals such as dispersal (Bargielowski et al., 2012; Evenden et al., 2014), resistance to diseases or chemical contamination (Townsend et al., 2010; Reid & Purcell, 2011), overwintering success (Sinclair, 2015) and fecundity (Arrese & Soulages, 2010).

Maintenance costs are assumed to stem from the cost of maintaining **structural size**, which is to say the amount of mass which cannot be used as a source of energy. Indeed, the **Absolute Energy Demand Hypothesis** (Calder, 1984) proposes that large animals require a larger absolute amount of energy reserves to maintain their larger size, which is diverted from fitness traits. This hypothesis therefore assumes that maintaining a larger body size is more costly than maintaining a smaller one. This is the reason why BCIs are constructed from several morphological measurements: typically, the measurement of a phenotypically plastic morphological trait which varies according to the intake of resources from the environment (e.g. mass), corrected for a measurement of the fixed structural body size (e.g. body length).

The use of BCIs is widely accepted in ecological studies involving vertebrates, where ethical concerns impair the ability to sacrifice animals to invasively assess the state of the resources present in their bodies (Stevenson et al., 2006; Warner et al., 2016). Their use in invertebrates, and especially insects, is far less widespread, where authors often rely on a simple measurement of size or mass (sometimes used interchangeably) to predict the amount of energy reserves experimental insects contain (Rolff & Joop, 2002; Amarillo-Suárez et al., 2011; Yoder et al., 2010; Sturm, 2016). The main assumption behind the use of body size instead of body condition is that the larger the insect, the higher the nutrient content (Berger et al., 2012). This assumption would be in line with the **Relative Efficiency Hypothesis**, which states that animals with a large body size have a more efficient energy use because mass specific metabolic rate decreases with body size (Kleiber, 1932; Thommen et al., 2019). Consequently, a large body size is advantageous compared to a smaller one (Tammaru et al., 2002).

A major limitation of this assumption is that insects undergo a complex life cycle in which the next developmental stage is achieved by shedding a rigid exoskeleton, the cuticle, therefore allowing further growth (Anderson, 1974; Reim, 2009; Knapp & Knappova, 2013, Moya-Laraño et al., 2008). In holometabolous insects, which undergo a complete metamorphosis between larval and adult stages, adult size is determined by the size reached by the larva before entering the pupal stage (Chafino et al., 2019). Therefore, the cuticle of the newly emerged insect is produced using resources of the previous developmental stage, which means that a simple measurement of structural size will represent past developmental conditions instead of current status (Anderson, 1974; Reim, 2009; Knapp & Knappova, 2013; reviewed in Knapp & Uhnavá, 2014), whereas a simple measurement of mass will include the mass of the exoskeleton, which can in some species be relatively heavy and costly to maintain (Locke, 1991). On the one hand, adult structural size reflects the amount of resources accumulated by the larvae and the absolute amount of resources available to the adult. On the other hand, the quantity of resources accumulated during adult life can vary and the relative amount of resources available for fitness traits may be constrained by structural size.

Insect body size has been shown to be positively correlated with fitness traits on numerous occasions (see Whitman, 2008, for review). Bigger females of the cricket *Teleogryllus commodus* lay more eggs (Sturm, 2016), as well as females of the moth *Streblote panda* (Calvo & Molina, 2005). Larger males sire a higher proportion of the offspring of females of the field cricket *Gryllus firmus* (Saleh et al., 2013). Body size is even assumed to be the main constraint on female fecundity (Honěk et al., 1993; Tammaru et al., 2002). On the other hand, studies which found a cost to a large structural body size in insects (reviewed in Blanckenhorn, 2000) identified mainly an increased predation risk but no direct energetic cost. Teuschl et al. (2007) observed a developmental cost of the larvae associated with the selection for a larger structural body size in the fly *Scathophaga stercoraria*, and pointed out that while there is strong empirical evidence of selection for large body size in insects, there is little evidence of the contrary.

Another important aspect to consider is that the use of BCIs in insects has been criticized because of their lack of correlation with the absolute amount of reserves present in an animal’s body and its fitness (see Wilder et al., 2016 for a review). Paradoxically, proponents of the use of BCIs argue that it is erroneous to invalidate BCIs on the basis of low correlation with absolute energy reserves since they are precisely designed to correlate with energy reserves corrected for a given size (i.e. relative energy reserves). Thus, their correlation with the absolute quantity of energetic reserves present in individuals is a sign that they do not properly control for structural size (DeBano, 2008; Peig and Green 2009, 2010).

We propose that the relevance of using BCIs in insects will depend on whether there is a significant cost to structural size. This cost would ultimately indicate the relative support for the Absolute Energy Demand or the Relative Efficiency Hypotheses in an organism in a given context. In this study we aimed to provide a thorough understanding of the biological consequences of interchangeably using body mass, structural size and body condition as a single proxy of energy reserves and predictor of fitness. We used an experimental approach using the mealworm beetle *Tenebrio molitor* (Coleoptera: Tenebrionidae), an insect massively reared as a source of food and for which breeder try to maximize body mass (Morales-Ramos et al., 2019; Costa et al., 2020). Fitness costs have been shown to increase in stressful environments (Schmid-Hempel and Schmid-Hempel 1998). We therefore assessed the cost of structural size on control and populations under food restriction at the larval and/or adult stage. First, we aimed to validate the use of structural size as a proxy of absolute energy reserves and BCIs as a proxy of relative energy reserves. This was done by sacrificing individuals, quantifying the lipids and sugars present in their bodies and correlating their absolute and relative values with body mass, structural size and BCIs. Second, we investigated the effect of body mass, structural size and body condition on the fitness parameter fecundity across feeding treatments.

## Materials and methods

Body mass and elytron size (as a measure of structural size) of adult female beetles were taken. From these measurements, we calculated 2 body condition indexes chosen from the literature, and explored their adequacy in controlling for the interdependence between a phenotypically plastic morphometric measurement and structural size. We did so on groups of female beetles having experienced food shortage or not at the larval and/or the adult stage (4 experimental treatments). From these females, a subset was killed after measuring in order to assess the quantity of energy reserves contained in their abdomens invasively, whereas a subset was allowed to reproduce in order to assess their fecundity. This way, we could evaluate the predictive power of body condition indexes on both the actual amount of reserves and female fecundity.

### Insect rearing and maintenance

Young larvae (∼ 1cm) were collected from a mass rearing outbred stock kept in the laboratory at the University of Burgundy in wheat bran (as a substrate, is also used as a source of food) with *ad libitum* access to water, and regularly supplemented with piglet flour (as a source of proteins) and a piece of apple (for carbohydrates and vitamins). Our insects are kept at 25°C and in the dark. After collection, we reared *T. molitor* larvae either in a rich or a poor larval feeding treatment, and subsequently maintained adults in a matching or mismatching environment, i.e. similar or dissimilar to the larval feeding treatment respectively.

### Generation of larval feeding treatments

After retrieving the larvae from the stock, we maintained them at a density of 250 larvae in a plastic box (L xH x l = 30 × 22 × 25 cm) in 5 litres of wheat bran with *ad libitum* access to water. We kept them in 2 separate containers according to the larval feeding treatment we subjected them to while controlling for density:

- In “**rich larval feeding**” conditions, larvae were kept similarly to our stock, that is to say in wheat bran + piglet flour + apple.
- In “**poor larval feeding**” conditions, larvae were simply kept in wheat bran with unlimited access to water and no further addition of piglet flour or apple. Importantly, larvae in this treatment took approximately 2 months longer to reach the pupal stage (data not shown).

All experimental individuals originate from these 2 boxes, which we checked daily for the presence of pupae.

The dynamics of pupation in our rearing conditions is such that a few larvae pupated early, preceding a “pupation peak” over 3 days, after which some larvae kept pupating late. Moreover, the growth of *T. molitor* is such that while getting closer to pupation, the food uptake of larvae stops as the insects are undergoing the physiological changes preparing metamorphosis (Connat et al., 1991). We retrieved pupae as soon as they were spotted, but we used in this experiment only the pupae formed during this “pupation peak”. This way, we assumed that we minimized as much as possible the effect of the removal of conspecifics on the accumulation of energy reserves by each individual larva. The resulting pupae were kept separately in a plastic box and checked daily for the presence of emerged adults in the teneral stage.

After emergence, adults were kept individually in Petri dishes supplied with wheat bran and a piece of apple for 5 days, time during which they reach sexual maturity. Five days after emergence the beetles were sclerotized enough to allow the determination of their sex by observation of their genitalia. As our study focused on variation in females body condition only, “poor larval” males were discarded, while “rich larval” males were transferred from their Petri dish into new plastic boxes where they were kept together according to their emergence date in wheat bran + *ad libitum* water + piglet flour + apple, waiting to be used in the later copulation experiments.

### Generation of adult female treatments

In this part of the experiment, wheat bran was replaced by thin flour in order to facilitate the recovery of the eggs by sieving (600 µm). Five days after emergence, females were maintained in Petri dishes according to the adult feeding treatment we subjected them to:

- “**poor adult feeding**” conditions, where females were kept in whole grain thin wheat flour, with *ad libitum* access to water.
- “**rich adult feeding**” conditions, where females were kept similarly but with addition of piglet flour and apple every second day.

The final sample sizes of the females of the four treatment combinations, aged 10 days old after emergence by the beginning of the experiment, were:

- 74 females in the “poor larval – poor adult” feeding treatment, 17 killed for assessment of their reserves, 56 allowed to reproduce
- 71 females in the “poor larval – rich adult” feeding treatment, 16 killed for assessment of their reserves, 55 allowed to reproduce
- 85 females in the “rich larval – poor adult” feeding treatment, 19 killed for assessment of their reserves, 54 allowed to reproduce
- 97 females in the “rich larval – rich adult” feeding treatment, 37 killed for assessment of their reserves, 56 allowed to reproduce

### Measures of body condition

#### Measure of morphological parameters

The mass of each female was measured on a Sartorius balance to a precision of 10^−5^ g before the first reproduction. Elytron length, a proxy of abdomen size (place of storage of reserves), of the *T. molitor* females was taken with a digital calliper (Mitutoyo, Absolute) to the nearest 0.01 mm. Repeatability of elytron length measurements was assessed by taking them twice on the same individual in a non-consecutive way. The females used for assessment of the energy reserves were weighed and measured before being frozen.

#### Body condition indexes calculations

For the calculation of BCIs, the correction by structural body size can be performed in different ways, resulting in several body condition indexes to chose from. The most frequently used in the literature for vertebrates (Blem 1990; Brown 1996; Schulte-Hostedde et al. 2005) and invertebrates (Jakob et al., 1996; DeBano, 2008; Kasumovic et al., 2009; Dubois et al., 2010) is the residuals from an Ordinary Least Squares regression of log transformed mass in relation to log transformed body size (referred to as OLS_resid_ hereafter; Jakob et al., 1996; Hayes and Shonkwiler, 2001). Its use has been criticized because despite controlling for the lack of independence between mass and structural size, it fails to control for the lack of independence between structural size and mass (see Green 2001; Peig and Green 2009, 2010; and supplementary material). As a consequence the use of the Standardized Mass Index (SMI) has been suggested as a good alternative (Peig and Green 2009; 2010; see supplementary materials, Figure S1). Peig and Green (2009) used the Thope-Lleonar model of scaling to estimate body condition, **using the slope from an standardized major axis regression of mass and structural size as the scaling exponent**

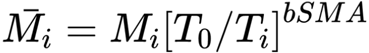

where *M*_*i*_ and *T*_*i*_ are the mass and structural size (elytron length) of an individual respectively. *T*_*0*_ is the population mean of the structural size, and bSMA is the slope of the regression between the log of *M*_*i*_ and the log of *T*_*i*_.

The benefit of Peig and Green’s (2009) formula is that it retains the original mass units. This results in a correction of our non-fixed morphological measure (i.e. mass) for structural size (i.e. elytron length). We refer to size-corrected morphological measures as “**scaled**”, i.e. **Scaled Mass Index (SMI)**. Since standardized major axis regressions standardise data, SMI can deal with Y and X variables measured on different scales. Its true error variance assumptions are also more realistic than ordinary least squares regressions in most cases (Warton et al., 2006; Green, 2001; see supplementary materials, Figure S1). We compare first the validity of these two BCIs based on their ability to predict both the relative amount of reserves present in our insect model system.

#### Quantification of the energy reserves

We used a dosage by colorimetry to assess the lipid content as well as the glycogen and the circulating carbohydrates present in the abdomen of the beetles, by using a protocol detailed in Rivero and Ferguson (2003). In brief, frozen beetles were dissected to remove their prothorax, head, elytra and legs. The remaining abdomens were homogenized in microcentrifuge tubes containing a 2% sodium sulfate solution and incubated in a 2:1 mix of chloroform:methanol solution for 24 hours. They were then centrifuged, to separate the pellet (containing glycogen) from the supernatant (containing fat and circulating sugars). The supernatant was divided into 2 equal parts. One part we used to quantify lipids with vanillin-phosphoric acid reagent. After a brief incubation, the optical density of the resulting solution was read in a spectrophotometer (SpectraMax®, Molecular Devices) at 525 nm and compared to a standard curve made of a serial dilution of sunflower seed oil. The other part was used to quantify mono and disaccharides from the supernatant and the pellet (after breaking down of the glycogen of the pellet), by incubating each of these fractions with anthrone reagent. The optical density of the resulting solutions was read at 625 nm and compared to a standard curve made of a serial dilution of glucose. As there was a strong correlation between circulating sugars and glycogen in the beetles (Pearson’s correlation of log-transformed sugars and glycogen [95% CI], *r* = 0.811 [0.725, 0.872], Figure S2), we considered both compounds together in the results.

Mono- and disaccharides from the supernatant represent circulating sugars mostly in the form of trehalose, as it is the main sugar present in the hemolymph of insects and serves as a storage for the glucose used in locomotory activity (Candy, 1989; Thompson, 2003), whereas glycogen stored in the fat body helps replenishing consumed trehalose (Candy, 1989). Diacylglycerol is another fuel used in locomotory activity, which is stored in the fat body (Canavoso, 2001; Williams & Robertson, 2008).

Similarly to the procedure applied to the morphological measures, we corrected the absolute quantities of lipids and sugars for structural size (i.e. elytron length) using Peig and Green’s (2009) formula. We refer to size-corrected body components as **scaled lipids** and **scaled sugars**.

A prerequisite to pooling data from different treatments when scaling metrics is that the relationship between log-transformed mass and log-transformed size does not change significantly according to our larval and adult feeding treatments (Peig & Green, 2009). This is the case in our dataset (standardized major axis regressions: larval feeding treatment: LRT = 2.558, *p-value* = 0.110; adult feeding treatment: LRT = 2.080, *p-value* = 0.149). Furthermore the relationship between all log-transformed energy reserves (lipids and sugars) and structural size also did not significantly differ between feeding treatments (standardized major axis regressions, all p-values > 0.05). Therefore we pooled all data together when calculating body condition indexes, scaled lipids and scaled sugars enabling comparisons between treatment groups.

### Copulations and fecundity assessment

Dnervich et al. (2001) showed that the mating of *T. molitor* females every second day prevented their sperm depletion. Our preliminary observations showed that females could lay eggs for a week following a single copulation event, and that when mated every second day they barely laid any eggs past 2 weeks after the first copulation. We thus designed our experiment in order to prevent any sperm depletion and capture most of the fitness of the females.

Ten days after emergence females were exposed to one male for 4 hours. During copulation, a male from the “rich larval” feeding treatment and a focal female were placed in an empty Petri dish containing only a piece of filter paper. We observed the pairs until we could record at least one copulation for each one of them, and we noticed that several events of copulation can take place in 4 hours. At the end of these 4 hours, the males were removed, put back together in a box and fed *ad libitum*, waiting to be reused 2 days later. Note that as males were not kept individually, they were randomly picked from their stock every second day, which reduces the chance that fecundity of the female would reflect the fitness of one male she would be paired with. Females on the other hand were individually placed in a new Petri dish supplemented with food according to their adult feeding treatment, and allowed to oviposit for 2 days until the next copulation event. The eggs of the previous Petri dish resulting from the previous reproduction event were sieved off of the flour and counted as a proxy for female fitness. There were in total 8 copulation sessions per female. Males that reached 17 days post emergence were removed from the experiment.

### Statistical analyses

All analyses were carried out in the statistical software R (R core team, 2021). Unless stated otherwise, we used the information-theoretic approach to achieve model selection (Burnham and Anderson, 2002). The relative strength of support of all candidate models was assessed via Akaike’s Information Criterion adjusted for small sample sizes (AICc) and AICc weights (ω), the adjusted *R*^*2*^ as defined by the MuMIn R package (Bartoń, 2020). We also report the effect size Cohen’s D for categorical variables and the partial *r* effect size for continuous variables (Nakagawa and Cuthill, 2007) and estimate the 95% confidence intervals by bootstrap (*n* = 10,000) using the “boot” R package (Canty and Ripley, 2021).

We first set out to validate whether morphometric measures and BCIs were reliable indicators of energy reserves. To that effect, we first tested the repeatability of the morphometric measure elytron length using the R package “rptR” and the strength of its correlation with mass (Stoffel and Nakagawa, 2017), both important preconditions of valid BCIs using these metrics (Peig and Green 2009; 2010). Second, we investigated Pearson correlations between measures of energy reserves and morphometric measures as well as with BCIs. Prior to correlations all morphometric measures and energetic reserves were log transformed.

Second, we aimed to test if absolute measures of energy reserves (using elytron length and body mass as proxies), relative energy reserves (using SMI as a proxy) or the combination of both were the best predictors of a fitness trait (i.e fecundity) in our insert model. With this aim in mind, we built five general linear models (GLM) with a negative binomial distribution, a log link function and with fecundity (i.e. number of eggs) as a response variable. Elytron length is fixed after emergence, it reflects body size independently of body condition, whilst SMI reflects body condition independently of structural size. We were able to include both elytron length and SMI in the same model because SMI and elytron were only weakly negatively correlated (Pearson’s correlation [95% confidence intervals]; *r* = -0.315 [-0.429; -0.191]) and the variance inflation factor was below 2, suggesting low collinearity when including both terms in a single model (Zuur et al. 2009). This allowed us to disentangle the relative support for the Relative Efficiency Hypothesis and the Absolute Energy Demand Hypothesis. In contrast, because of collinearity between elytron length and mass both parameters were always assessed in separate models. Thus, the five different models assessed were: fecundity according to body mass, fecundity according to structural size (elytron length), fecundity according to body condition (SMI), fecundity according to both structural size and body condition and the null model.

Third, we predicted that food limitation at the larval and/or the adult stage will accentuate a potential cost to structural size and thus will change the relationship between fitness and body condition. We investigated if food restriction resulted in a different relationship between fecundity and elytron length/body mass/SMI using a GLM with a negative binomial distribution, a log link function and with fecundity (i.e. number of eggs) as a response variable. We build sixteen models with the explanatory variables elytron length, body mass, SMI and interactions between feeding treatment and elytron length/body mass/SMI. Once again, the effect of mass was assessed in separate models from elytron length and SMI due to collinearity.

Since selection and/or environmental plasticity at the larval stage can result in differences in morphology of adults at pupal emergence (Teuschl, Reim & Blanckenhorn, 2006), we tested whether larval feeding treatment affected morphology and condition at emergence from pupation. We therefore assessed differences in mean elytron size, mass and SMI between larval treatments, after checking for homogeneity of variance using the Fligner-Killeen test. We then assessed mean differences between larval treatment groups using linear models for elytron length and mass and a general least square (GLS) model for SMI to control for significant heterogeneity (see results).

## Results

### 1. Validation of structural size and body condition indexes as measure of absolute and relative energy reserves respectively

Our measures of elytron length were significantly repeatable (p < 0.05) with an *R* [95%CI] of 0.921 [0.902, 0.936] and were strongly correlated with mass (Pearson’s correlation [95% confidence intervals]; *r* = 0.776 [0.729; 0.816]). Our results confirm that elytron length is an ideal indicator of structural size in our insect given its high repeatability in measurement and its strong correlation with mass. Moreover, its size is fixed after hatching from the pupal stage, since the cuticle gets tanned and sclerotized after emergence (i.e. it does not vary during the life of the insect)

Both elytron length and mass were highly correlated with absolute values of lipids and sugars (**Table 1**). In contrast, mass correlated poorly with the amounts of lipids and sugars that were scaled for structural size (**Table 1**). OLS_resid_ and SMI on the other hand significantly correlated with quantities of lipids and sugars which were scaled for body size (**Table 1**). However, importantly, in contrast with SMI, OLS_resid_ additionally correlated significantly with absolute values of lipids and sugars (**Table 1**), consistent with the fact that OLS_resid_ is correlated with structural body size (see supplementary material, Figure S1, for more details). Since BCIs have been introduced within the context of the Absolute Energy Demand Hypothesis, they should not be expected to correlate with the absolute amount of reserves, but with the amount of reserves relative to structural size instead (DeBano, 2008; Peig & Green 2009; 2010). Thus our results confirm that OLS_resid_ is a biased index of body condition since it does not control adequately for the codependence between structural size and mass, whereas SMI does. We therefore excluded OLS_resid_ from all subsequent analyses, and focus on elytron length, mass and SMI.

**Table 1:** Pearson’s correlation between log-transformed components of body condition and log-transformed body measures or body condition indices.

|  | *log(Lipids)* | *log(Glucose+glycogen)* | *log(Scaled lipids)* | *log(Scaled glucose+glycogen)* |
| --- | --- | --- | --- | --- |
| *a. Body measures* |  |  |  |  |
| *log(Elytron length)* | *r* = 0.612<br>(0.462; 0.727) | *r* = 0.703<br>(0.580; 0.795) | N/A | N/A |
| *log(Mass)* | *r* = 0.685<br>(0.556; 0.782) | *r* = 0.740<br>(0.628; 0.821) | *r* = -0.144<br>(-0.342; 0.066) | *r* = -0.093<br>(-0.295; 0.118) |
| *b. Body condition indices* |  |  |  |  |
| *OLS residuals* | *r* = 0.374<br>(0.180; 0.541) | *r* = 0.350<br>(0.153; 0.520) | *r* = 0.326<br>(0.126; 0.500) | *r* = 0.341<br>(0.143; 0.513) |
| *Scaled mass index (SMI)* | *r* = 0.169<br>(-0.041; 0.364) | *r* = 0.097<br>(-0.113; 0.299) | *r* = 0.452<br>(0.269; 0.603) | *r* = 0.424<br>(0.236; 0.581) |

### 2. The relevance of structural size and body condition as predictors of fitness in an insect model

In this section, we assessed five different models of female fecundity: fecundity according to body mass, fecundity according to structural size (elytron length), fecundity according to body condition (SMI), fecundity according to both structural size and body condition and the null model. The latter was the model which was the best supported (**Table 1**; Δ**AICc *=*** 2.066; *ω =* 0.622; R^2^ = 0.055), with both predictors positively associated with fecundity (**Figure 1**; elytron length: partial-*r* [95%CI] *=* 0.156 [0.020; 0.277]; SMI: partial*-r* [95%CI] *=* 0.223 [0.109; 0.326]). Among the other models, when predictors were assessed individually, models with body mass or SMI had stronger support than the null model (**Table 2**; **Figure 1**; body mass: R^2^ = 0.037; partial-*r* [95%CI] *=* 0.197 [0.075; 0.315] and SMI: R^2^ = 0.033; partial-*r* [95%CI] *=* 0.183 [0.071; 0.291]) but interestingly this was not the case for elytron length (**Table 2**; **Figure 2**; R^2^ = 0.007; partial-*r* [95%CI] *=* 0.086 [-0.044; 0.212]). Therefore the effect of body size on fecundity was only apparent and statistically supported when accounting for body condition (**Figure 1**), and the effect of body condition was stronger and better supported when accounting for body size.

**Figure 1:**
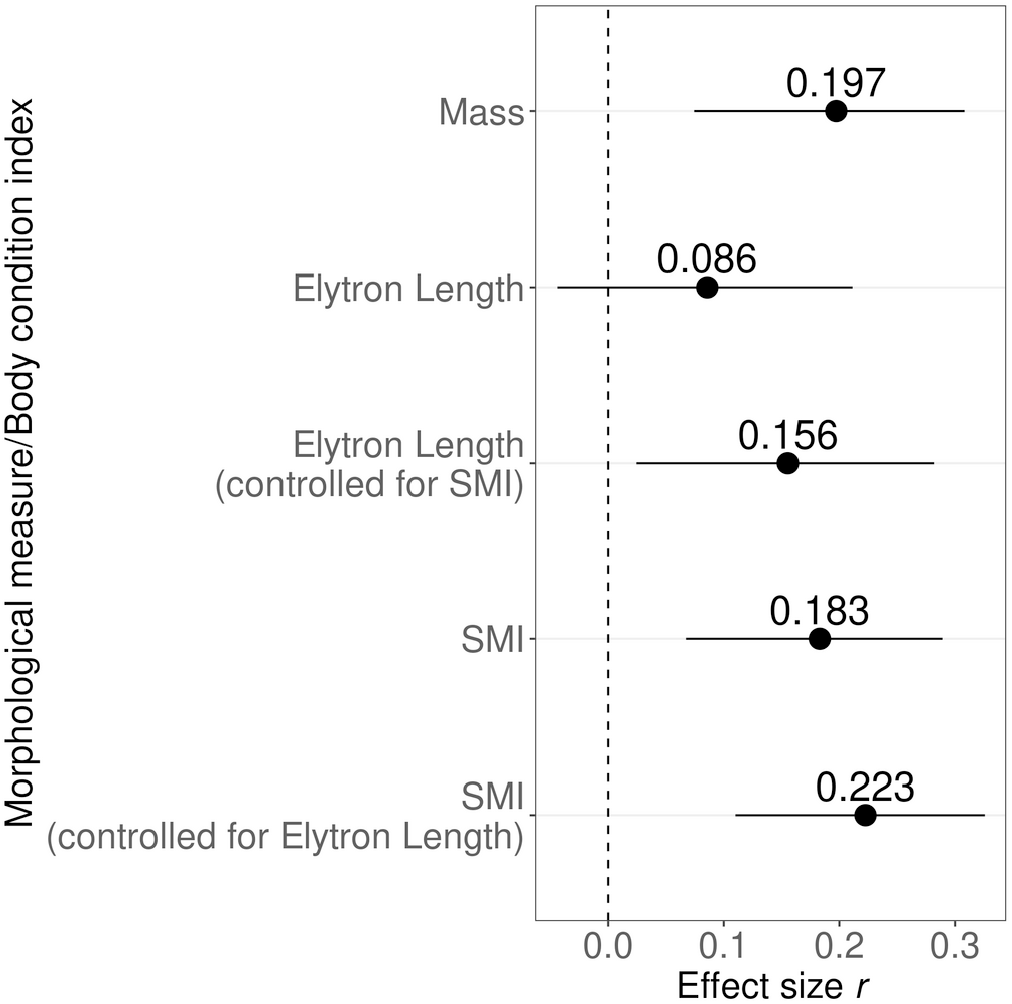
Effect size of the effect of mass (model 2; Table 2), elytron length (model 5; Table 2), elytron length when controlling for SMI (model 1; Table 2), SMI (model 3; Table 2) and SMI controlling for elytron length (model 1; Table 2) on total fecundity.

**Figure 2:**
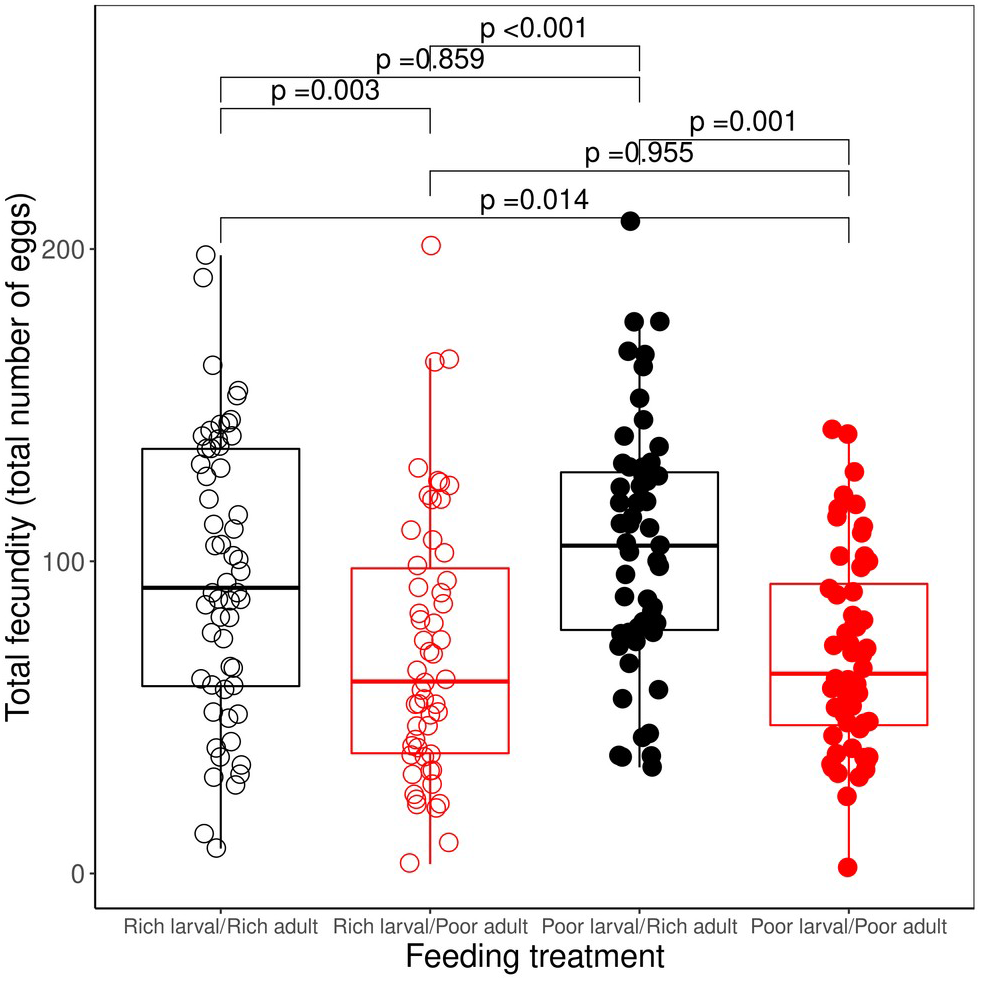
Boxplot of total fecundity according to larval and adult feeding treatments. Each data represents a value from a female. Also shown are p-values from a Tukey’s honest significance test comparing the effect of different levels feeding treatments on fecundity (controlling for elytron length and SMI) according to model 3 in Table 3 (the most parsimonious model without interaction terms). Feeding restriction had a significant effect on fecundity at the adult stage but not at the larval stage.

**Table 2:** Five candidate models of the effect of elytron length, body mass and SMI on fecundity according to a general linear model (GLM) with negative binomial distribution. Model rank, the model structure, model degrees of freedom (df), model log-likelihood (LogLik), model Aikaike information criterion for small sizes (AICc), AICc weight (*ω*) and model R^2^ are shown.

| *Model Rank* | *Model* | *d f* | *LogLik* | *AICc* | $\Delta AICc$ | $\omega$ | $R^2$ |
| --- | --- | --- | --- | --- | --- | --- | --- |
| 1 | *Fecundity ~ Elytron length + SMI* | 4 | -1133.662 | 2275.509 | 0.000 | 0.622 | 0.055 |
| 2 | *Fecundity ~ body mass* | 3 | -1135.732 | 2277.575 | 2.066 | 0.221 | 0.037 |
| 3 | *Fecundity ~ SMI* | 3 | -1136.192 | 2278.494 | 2.985 | 0.140 | 0.033 |
| 4 | *Null model* | 2 | -1139.856 | 2283.767 | 8.258 | 0.010 | 0.000 |
| 5 | *Fecundity ~ Elytron length* | 3 | -1139.106 | 2284.322 | 8.813 | 0.008 | 0.007 |

### 3. Is the cost of size on fitness dependent on resource availability during the larval and adult stages?

In this next section, we investigated whether the effect of food limitation at the larval and/or adult stage changes the relationship between fecundity and elytron length/mass/body condition. We expected that food limitation at the larval and/or the adult stage would accentuate a potential cost to structural size and we therefore expected statistical support for the effect of an interaction between body condition and feeding treatment on fecundity. Feeding treatment was an important predictor of fecundity (ΔAICc = 23.588; Table 3; Figure 2) and, as predicted, the effect of an interaction between food treatment and SMI was supported by model selection (ΔAICc = 3.107; Table 3; Figure 3). None of the other interactions were supported by model selection (ΔAICc < 2; Table 3). Therefore, the effect of SMI on female fecundity was dependent of the feeding treatment, but not mass or elytron length.

**Table 3:** Sixteen candidate models of the effect of elytron length, body mass, SMI and interactions between feeding treatment and elytron length/body mass/SMI on fecundity according to a general linear model (GLM) with negative binomial distribution. The effect of mass was assessed separately from elytron length and SMI due to collinearity. Models with interaction terms included the corresponding additive terms, but for simplicity the shortcut formula is presented (i.e. a model with y ∼ x + z + x:z is presented as y ∼ x * z). Model rank, the model structure, model degrees of freedom (df), model log-likelihood (LogLik), model aikaike information criterion for small sizes (AICc), AICc weight (*ω*) and model coefficient of determination (R^2^) are shown.

| **Model Rank** | **Model** | **df** | **LogLik** | **AICc** | **$\Delta AICc$** | **$\omega$** | **$R^2$** |
| --- | --- | --- | --- | --- | --- | --- | --- |
| 1 | *Fecundity ~ feeding treatment \* SMI + elytron length* | 10 | -1115.436 | 2251.920 | 0.000 | 0.615 | 0.198 |
| 2 | *Fecundity ~ feeding treatment \* elytron length + feeding treatment \* SMI* | 13 | -1113.635 | 2255.028 | 3.107 | 0.130 | 0.211 |
| 3 | *Fecundity ~ feeding treatment + elytron length + SMI* | 7 | -1120.437 | 2255.401 | 3.480 | 0.108 | 0.161 |
| 4 | *Fecundity ~ feeding treatment \* elytron length + SMI* | 10 | -1117.702 | 2256.453 | 4.532 | 0.064 | 0.182 |
| 5 | *Fecundity ~ feeding treatment \* SMI* | 9 | -1119.304 | 2257.460 | 5.540 | 0.039 | 0.170 |
| 6 | *Fecundity ~ feeding treatment + mass* | 6 | -1122.749 | 2257.892 | 5.971 | 0.031 | 0.143 |
| 7 | *Fecundity ~ feeding treatment + SMI* | 6 | -1124.001 | 2260.395 | 8.474 | 0.009 | 0.134 |
| 8 | *Fecundity ~ feeding treatment \* mass* | 9 | -1121.511 | 2261.876 | 9.955 | 0.004 | 0.153 |
| 9 | *Fecundity ~ feeding treatment + elytron length* | 6 | -1126.790 | 2265.973 | 14.053 | 0.001 | 0.112 |
| 10 | *Fecundity ~ feeding treatment* | 5 | -1128.006 | 2266.290 | 14.370 | <0.001 | 0.102 |
| 11 | *Fecundity ~ feeding treatment \* elytron length* | 9 | -1124.490 | 2267.833 | 15.912 | <0.001 | 0.130 |
| 12 | *Fecundity ~ elytron length + SMI* | 4 | -1133.662 | 2275.509 | 23.588 | <0.001 | 0.055 |
| 13 | *Fecundity ~ mass* | 3 | -1135.732 | 2277.575 | 25.655 | <0.001 | 0.037 |
| 14 | *Fecundity ~ SMI* | 3 | -1136.192 | 2278.494 | 26.574 | <0.001 | 0.033 |
| 15 | *Null model* | 2 | -1139.856 | 2283.767 | 31.846 | <0.001 | 0.000 |
| 16 | *Fecundity ~ elytron length* | 3 | -1139.106 | 2284.322 | 32.402 | <0.001 | 0.007 |

**Figure 3:**
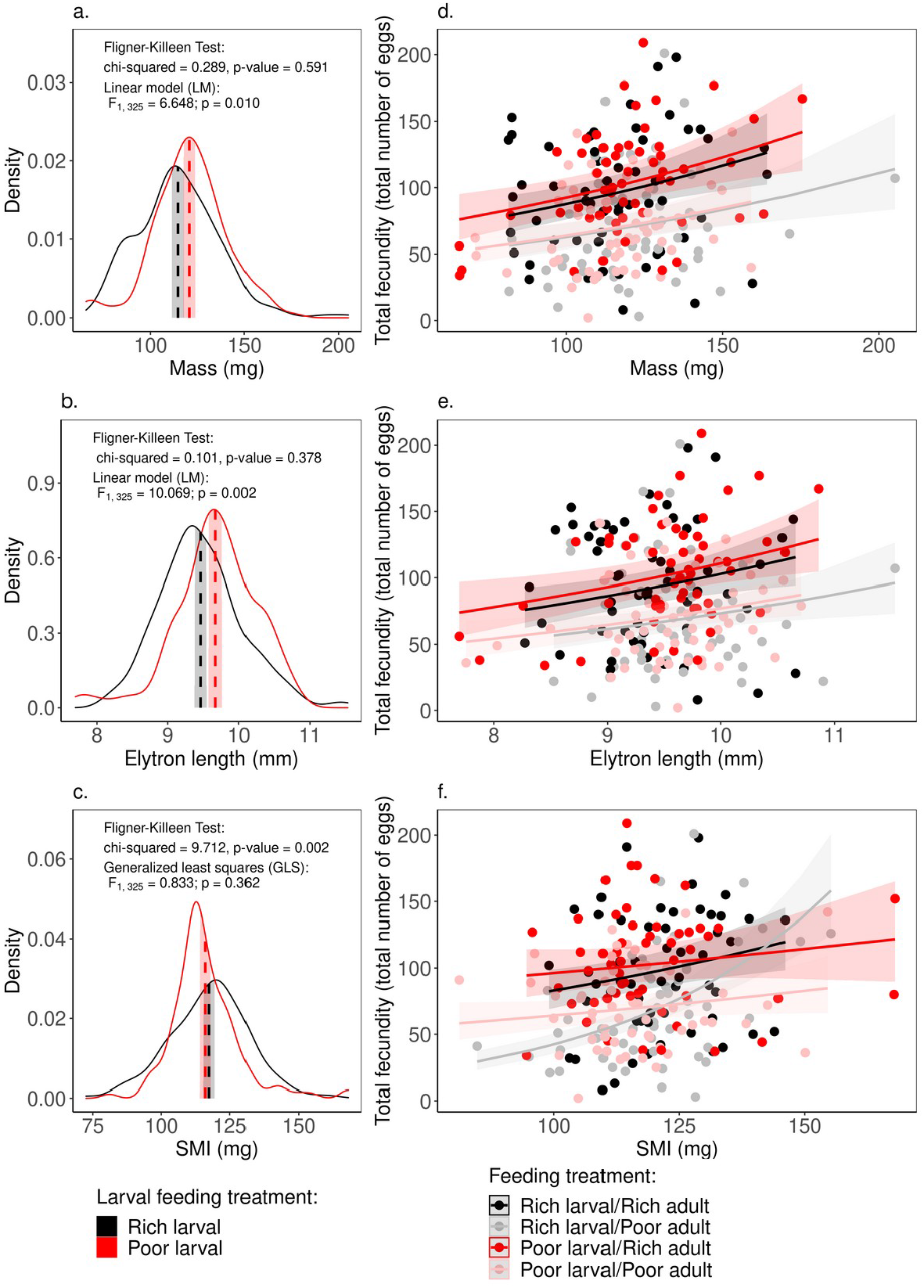
Density curves elytron length (a.), mass (b.) and SMI (c.) according to the larval feeding treatment. The mean and 95%CI intervals are shown as a dashed line along and shaded ribbons. Also shown are the output from the Fligner-Killeen Test of Homogeneity of Variances to test differences in variation between larval feeding treatment groups. Finally differences in mean between larval treatment groups were tested using linear models (LM) for mass (a.) and elytron length (b.) and whilst general lest squares (GLS) were used for SMI to control for heterogeneity between larval feeding groups. The effect of mass (d.), elytron length (e.), and SMI (f.) on fecundity according to feeding treatment are plotted with fitted lines and 95% confidence intervals.

Contrary to our expectations, the restriction of food at the larval stage (in “poor larval-poor adult” and “poor larval-rich adult”) did not increase the effect of SMI on fecundity. Instead, we found that the positive effect of SMI on fecundity was large and at its strongest within the rich larval and poor adult feeding groups (“rich larval-poor adult”: partial *r* [95%CI] = 0.476 [0.268; 0.632]), whilst the effect was small and not statistically supported in the remaining feeding treatments (in decreasing order: “poor larval-rich adult”: partial *r* [95%CI] = 0.206 [-0.114; 0.490]; “rich larval-rich adult”: partial *r* [95%CI] = 0.148 [-0.125; 0.367]; “poor larval-poor adult”: partial *r* [95%CI] = 0.126 [-0.104; 0.366]). This result can best be explained by investigating the variation in elytron length, mass and SMI within larval feeding treatment groups. Indeed, individuals originating from the poor larval feeding treatment emerged as adults with larger structural size (Cohen’s d = 0.354 [0.125-0.583]; **Figure 3a**.) and heavier mass (Cohen’s d = 0.287 [0.061-0.514]; **Figure 3b**.) on average than individuals originating from the rich larval feeding treatment. In contrast, adults originating from both treatments have a similar body condition (SMI: Cohen’s d = 0.100 [-0.018; 0.416]; **Figure 3c**.). However, the variability in SMI differed between the larval feeding treatments, adults originating from the poor larval feeding treatment showing less dispersion in the SMI scores than adults originating from the rich larval feeding treatment (Fligner-Killeen test: *χ*^*2*^ = 9.712; *p-value* = 0.002; **Figure 3c**.). We do not observe this heterogeneity between larval feeding treatments on either structural size (elytron length) or mass (**Figure 3a. and b**.). It therefore appears that individuals form the poor larval food treatment emerged larger and more individuals were in peak body condition (lower variation in SMI around the mean) compared to the rich larval treatment. This reduced variation in SMI explains the weak effect of SMI on fecundity within adults originating from the poor larval feeding treatment (**Figure 3c. and e**.).

The fact that the effect of SMI was stronger in rich larval-poor adult feeding treatment compared to rich larval-rich adult feeding treatment is in line with our prediction that food restriction would accentuate the effect of body condition on fecundity. Further evidence of the stressful effect of food restriction on our animals comes from the fact that we also observed a higher mortality in females in the rich larval and poor adult feeding groups, whilst there was no mortality in the poor larval and rich adult feeding group (ΔAICc = 14.308; mortality: Rich larval/Poor adult = 18%; Rich larval/Rich adult = 7%; Poor larval/Poor adult = 2%; Poor larval/Rich adult = 0%). We found no support of effects of SMI, mass, elytron length or interactions on mortality (ΔAICc < 2; Table S1; Figure S3).

## Discussion

Many invertebrate studies use mass, size and body condition interchangeably as a single proxy of energy reserves and predictor of fitness (Amarillo-Suárez et al., 2011; Yoder et al., 2010; Sturm, 2016). Moreover, in both vertebrate and invertebrate studies, body condition indexes are frequently “invalidated” on the basis that they correlate poorly with absolute measures of energy reserves (Schulte-Hostedde et al. 2001; Labocha, Schutz & Hayes 2014; Wilder et al., 2016; McGuire et al. 2018). With this study, we aimed at shedding light on whether the concept of body condition, and thus the use of BCIs, is relevant in insects, or if on the contrary, absolute measures of energy reserves are more relevant. Indeed, in insects there is extensive evidence of a benefit to a larger structural size, but no evidence of a direct cost. These costs or benefits would ultimately indicate the relative support for the Absolute Energy Demand or the Relative Efficiency Hypotheses in an organism in a given context.

From a methodological point of view, our results confirm that simple measurements of size and mass are a good measure of the absolute amount of lipids and sugars present in *T. molitor* females. Regarding BCIs on the other hand, the correlation between SMI and the absolute amount of reserves was weak and not statistically supported. The fact that a BCI does not correlate or only weakly to an absolute amount of reserves has been criticized by several authors (Labocha, Schutz & Hayes 2014; Wilder et al., 2016). It is important to note however that since BCIs have been introduced within the context of the Absolute Energy Demand Hypothesis, they are supposed to account for energy expenditure relative to body size. Thus, they are not expected to correlate with the absolute amount of reserves, but with the amount of reserves relative to structural size instead (DeBano, 2008; Peig & Green 2009; 2010). In this case, it is precisely the strong correlation between SMI and relative energy reserves and the lack of correlation with absolute energy reserves that validates BCI as a good indicator of body condition.

In vertebrates, prior to Peig and Green’s (2019) study, OLS_resid_ was the most prevalent BCI in use but has since largely been displaced by SMI as a consequence of OLS_resid_ biases in estimating body condition (Peig & Green 2009; 2010). Our study expands the findings of the biases of OLS_resid_ to an insect model, where OLS_resid_ do not appropriately control for the dependence of body mass on body structural size, and consequently correlates with both absolute and relative body reserves. Furthermore, SMI are likely to be a superior to OLS_resid_ in most systems since the dependence of mass on structural size will violate OLS error assumptions in most scenarios (DeBano, 2008; Peig & Green 2009, 2010; and, see supplementary material, Figure S1). Therefore, we converge with several other authors who advise on BCIs which are based on standardised major axis regressions rather than ordinary least squares in any ecological study (Kotiaho, 1999, Green, 2001; Peig & Green, 2009, 2010).

From an ecological point of view, models with both elytron length and body condition were better supported than models with a simple measurement of mass in *T. molitor* females. Furthermore, the effect of body size on fitness was only discernable when controlling for the effects of body condition. Thus, our results show that after accounting for the cost of body size maintenance (consistent with the Absolute Energy Demand Hypothesis), larger individuals have a higher fitness (consistent with the Relative Efficiency Hypothesis), contradicting previous observations in several insect species that structural size does not represent a significant cost (Tammaru et al., 2002; Calvo & Molina, 2005; Honěk et al., 1993; Wilder et al., 2016).

As predicted, the observed effect of the interaction between SMI and feeding treatment indicates that the relative weight of the Relative Efficiency Hypothesis and Absolute Energy Demand Hypothesis is dependent on environmental resource availability. An examination of the effects of larval feeding treatments on the morphological parameters reveals that adult females originating from the poor larval feeding treatment show a similar average but a reduced variance in their body condition compared to females originating from the rich larval feeding treatment. This reduced variance is likely to be responsible for the weak relationship between SMI and fecundity in females which went through food restriction at the larval stage. In contrast, the higher variation in body condition in females that had unrestricted access to food at the larval stage resulted in a strong and positive relationship between body condition and fitness but only when environmental food access was limited at the adult stage.

Unlike body condition, structural size and mass show a divergence in their mean values between larval feeding treatments but a similar dispersion: females originating from the poor larval feeding treatment reach a larger structural size than their well-fed counterparts. Our experimental design does not allow us to conclude whether the reduced variance in body condition and increased body size in the poor larval treatments is the result of developmental plasticity or differential mortality. On the one hand, although we did not find a higher larval mortality between our feeding treatments, we did not monitor mortality during metamorphosis. As a very costly life-history event which lives insects vulnerable to predation, or, in our setting, cannibalism, we could expect it to be an important culling point in *T. molitor*, as well as numerous insect species (Ichikawa & Kurauchi, 2009; Lowe et al., 2021). On the other hand, larvae reared in the poor larval feeding treatment took approximately two months extra to reach the pupation stage compared to larvae reared in the rich larval feeding treatment, which indicates a certain degree of developmental plasticity. *T. molitor* is indeed known to be very plastic in its larval developmental time depending on access to food among other parameters (Cotton & St. George, 1929).

It has been shown in several insect species that restricted access to food at the larval stage can sometimes lead to a delayed maturation at a smaller adult body size (Day & Rowe, 2002, Dmitriew et al., 2009). We observed the contrary, that adult females emerging from the poor larval feeding treatment were slightly bigger (2 % difference in the elytron size) than females emerging from the rich larval feeding treatment. It is therefore likely that this reduced variance in body condition is the result of both developmental plasticity and differential mortality prior to adult emergence. Moreover, we found higher adult mortality for insects well fed at the larval stage but food restricted at the adult stage, independently of body condition. Thus, selection and/or developmental plasticity which resulted from the restriction of food at the larval stage resulted in insects which were better able to cope with food restriction at the adult stage, regardless of their starting body condition. In contrast, emergence at suboptimal body condition for females with unrestricted access to food at the larval stage resulted in reduced fitness when food was restricted at the adult stage. Thus, the potential adaptive benefit of emerging earlier in favourable environmental conditions comes at a fitness cost for females with low body condition if the environment changes unfavourably at the adult stage.

The reduced variance in SMI but divergence in mean structural size and body mass indicates that the amount of resources relative to structural size, instead of the absolute amount of reserves, is optimized prior to adult emergence in *T. molitor* females. This means that there is a cost to structural size in our insect model, consistent with the Absolute Energy Demand Hypothesis. At the same time, the larger structural body size in females originating from the poor larval feeding treatment could be the result of a maximisation of the mass-specific metabolic rate during the larval development under food restriction, which is consistent with the Relative Efficiency Hypothesis. Considering that our insect model is a flightless burrowing insect, we can expect the cost of structural size due to locomotion is reduced compared to other insect species in which flight is an additional source of costs (Dixon & Kindlmann, 1999; Williams & Robertson, 2008).

Therefore, we urge authors to take into account body condition in the form of BCIs in insect ecological studies in order to highlight fitness effects which might have been hidden by the use of structural size only or a measure which is not independent from it. From a more applied perspective, our study might help insect breeders maximize nutrient content of the insects, by maximizing their body condition, over the production of components of structural size (such as the cuticle) which have a poor availability to the vertebrate digestive system. In contrast, the production of insect by-products such as chitin, the main component of the insect cuticle, and thus, of insect structural size, might be improved by maximizing structural size over body condition.

To conclude, our study demonstrates that the concept of body condition, and thus correcting the amount of energy reserves for structural body size in invertebrates, is more relevant than using a simple measurement of mass as a proxy for the amount of reserves available for the allocation to fitness traits (Knapp & Knappova, 2013; Kasumovic & Andrade, 2006; Kasumovic et al., 2009; Moya Laraño et al., 2008). Our analyses show that this does not stem from stronger support of the Absolute Energy Demand Hypothesis over the Relative Efficiency Hypothesis in our insect model. Instead, the use of an appropriate BCI allowed us to detect both a cost (since higher body condition predicts fecundity) and a benefit to structural body size (since larger females have a higher fitness at a given body condition) in our insect model. This confirms, as stated by (Reim et al. 2006), that both the Absolute Energy Demand Hypothesis and the Relative Efficiency Hypothesis are not mutually exclusive, and instead that the fitness of an individual will be the result of their combined action. Applying a similar approach in more studies on various animal models might reveal the relative importance of both these processes in different ecological contexts.

## Acknowledgements

We thank Maria Texeira Brandao for the help with the quantification of sugars and lipids. We thank Coralie Dal & Aude Balourdet for experimental support. We thank Morgan David for initial discussion, and Jens Rolff for comments on the manuscript. This work was funded by ANR-JCJC-0134 & ANR-14-CE02-0009 to YM, and the DFG grant 5026 FOR InsectInfect.

## Authors contributions

Conceived the study and designed the experiments: CZ, YM & MG. Conducted the experiments: CZ & MG. Analyzed the data: MG. Wrote the manuscript: CZ & MG with contributions from YM.

## Data archiving statement

the data will be made available on Dryad upon acceptance for publication.

