## Supplemental information for "Size matters, so does condition: the use of a body condition index reveals the costs and benefits of structural body size in an insect"

### Supplementary Figures

##### Figure S1: Relationship between mass and elytron size according to OLS, SMA and MA regressions.

OLS regressions assume that the dependent variable X is independent of the response variable Y and that X is free of errors. Both assumptions are violated since structural size will tend to be greater in individuals of higher condition and X will suffer from some error. It is also important to note that measurement is not necessarily the main source of error in structural size, a larger source of error may be variation between individuals in how well measures of structural size actually reflect true body size due to variation in body shape (Warton et al., 2006, Green, 2001). The result of these violations in assumptions is that the slope of the OLS regression will be underestimated, resulting in an over-estimation of condition for large individuals and vice-versa for smaller individuals. In our study, although by construction $OLS_{resid}$ is not correlated with structural size (**Figure S1c.**), it is positively correlated with mass (**Figure S1f.**). Therefore, as predicted, the slope of the OLS regression between the log of mass and the log of elytron size is underestimated, resulting in an over-estimation of condition for large individuals and vice-versa for smaller individuals (**Figure S1a**).

Unlike OLS regressions, major axis (MA) regressions assume that X is not free of errors, that the errors in X and Y are interdependent, and that the ratio in error variance in Y and X is equal to 1 (Green, 2001; Warton et al., 2006). However, the error variance is expected to be greater in measurements of plastic phenotypic traits (such as mass) than in measurements of structural size (Rising and Somers 1989; Green, 2001). In addition, measurements of plastic phenotypic traits and structural size are generally not measured within comparable scales. Therefore, in most cases, the error variance ratio Y/X assumption of MA regression will be violated and will lead to inflated slopes that will approach an OLS regression slope of structural size (i.e. X) on the plastic phenotypic trait (i.e. Y) (Green, 2001). In our study residuals from major axis (MA) regressions did not correlate with mass (**Figure 1e.**) but strongly negatively correlated with structural size (**Figure**

**1h.**). Therefore, when applying an MA regression to log-transformed mass to log-transformed elytron size, we found as stated by Green (2001) that the slope was much steeper than the slope of all other regression methods and is overestimated.

To address the interdependence between mass and size, the use of the residuals from standardised major axis (SMA) regressions (otherwise known as reduced major axis regression) have been proposed to be better suited for the calculation of BCIs than OLS or MA regressions (Green 2001, Peig and Green 2009, 2010). SMA regressions assume that the ratio of error variance Y/X is equal to the ratio of the true variance Y/X. Since SMA regressions standardise data, it can deal with Y and X variables measured on different scales. Its true and error variance assumptions are also more realistic than both OLS and MA regressions in most cases (Warton et al., 2006, Green, 2001). When applying the SMA regression to our data, the slope of the SMA regression was within the slopes of OLS and MA regression methods (**Figure S1a.**). However SMA regression does not completely correct for the dependence between mass and size since when plotting SMA residuals against mass and elytron length, they weakly correlated positively with the former (**Figure S1d.**) and negatively with the latter (**Figure S1c.**). Because the slope of the SMA regression is used as the scaling exponent, it is perfectly correlated with log-transformed SMA residuals (**Figure S1b.**).

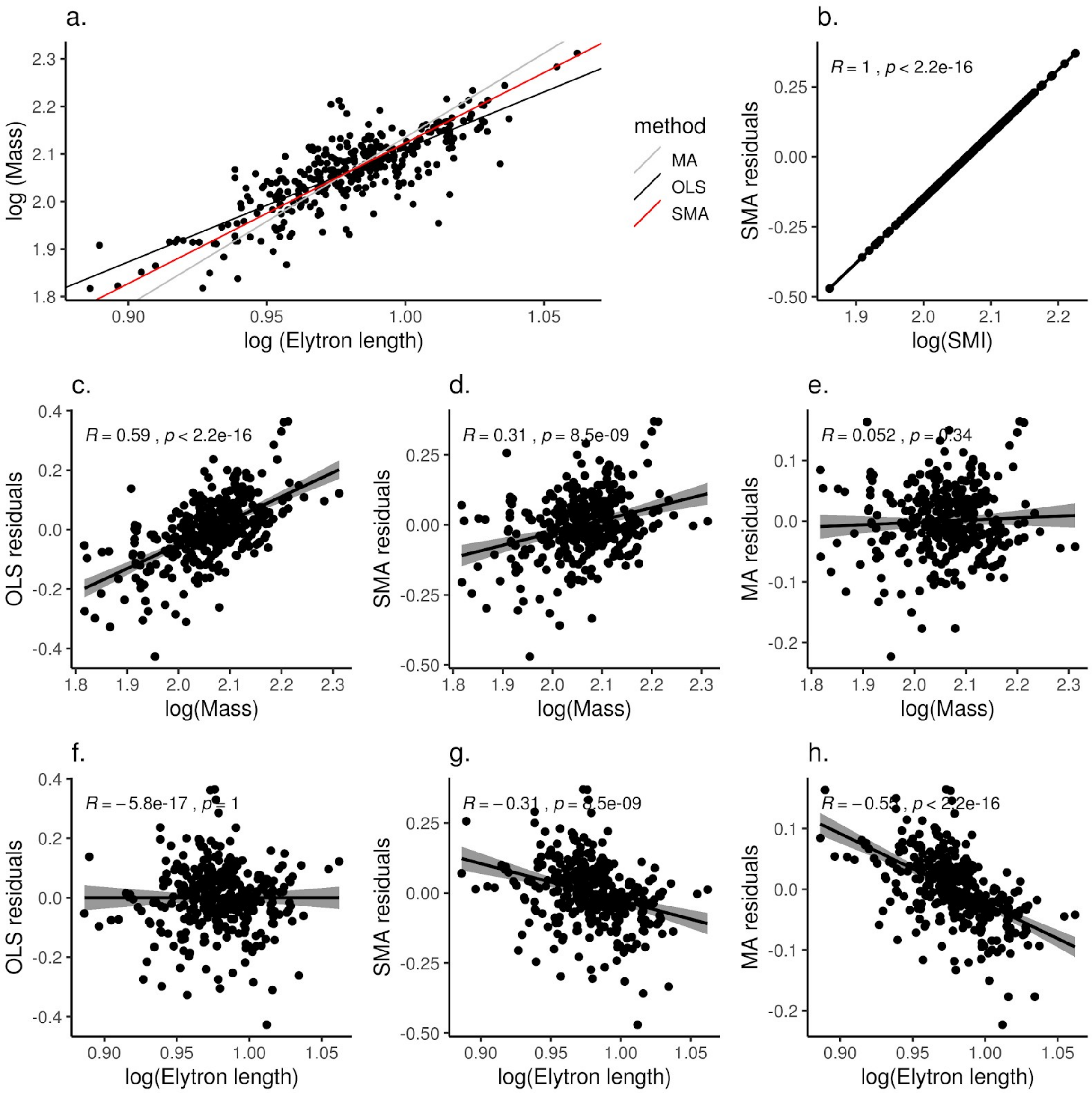


**Figure S2**: a. Ordinary Least Squares (OLS, in black), Major Axis (MA in grey), and Standardized Major Axis (SMA, in red) regressions between the logarithm of elytron length and the logarithm of mass of individual *Tenebrio molitor* females. The correlation between the residuals of the OLS regression mass and elytron length are represented respectively in c. and f.. The correlation between the residuals of the SMA regression, mass and elytron length are represented in d. and g., and the correlation between the residuals of the MA regression, mass and elytron length are represented in e. and h.. b. Correlation between the residuals of the SMA regression and the logarithm of the Standardized Mass Index (SMI) calculated according to the formula of Peig & Green (2009). Each dot represents an experimental individual.

**Figure S2: Correlation between glucose and glycogen quantities in the bodies of *Tenebrio molitor* females.**

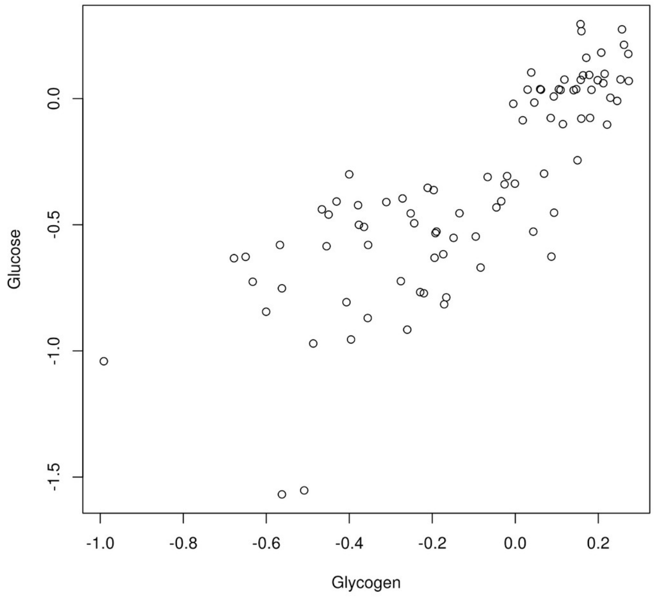


**Figure S2**: Correlation between the logarithm of glucose and glycogen quantities in the subset of *T. molitor* females killed at the start of the experiment. Each dot represents an individual.

**Figure S3: Proportion of dead *T. molitor* females by the end of the experiment according to the larval and adult feeding treatments.**

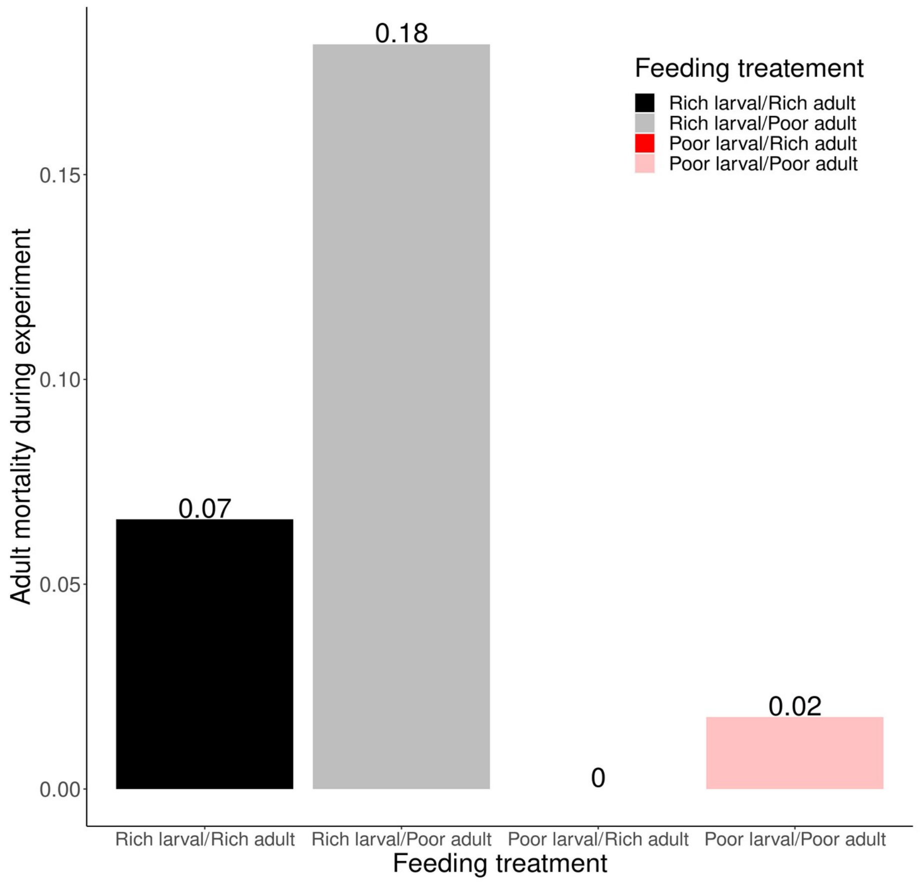


**Figure S3**: The proportion of females that died during the course of the experiment according to larval and adult feeding treatments

We hypothesised that there might be differential adult mortality between different feeding treatments and according to morphological measures and body condition. To test if food restriction resulted in a difference in mortality during the course of the experiment, we examined differential mortality according to elytron length, body mass, SMI and interactions between feeding treatment and elytron length/body mass/SMI on adult mortality during the course of the experiment using a GLM with binomial distribution. The effect of mass was assessed separately from elytron length and SMI due to collinearity.

Evidence of the stressful effect of food restriction on our animals comes from the fact that we observed a higher mortality in females in the rich larval and poor adult feeding groups, whilst there was no mortality in the poor larval and rich adult feeding group (ΔAICc = 14.308; mortality: Rich larval/Poor adult = 18%; Rich larval/Rich adult = 7%; Poor larval/Poor adult = 2%; Poor larval/Rich adult = 0%; Figure S3; Table S2). We found no support of effects of SMI, mass and elytron length or interactions on mortality (ΔAICc < 2; Table S1).

**Table S1:** Sixteen candidate models of the effect of elytron length, body mass, SMI and interactions between feeding treatment and elytron length/body mass/SMI on mortality according to a general linear model (GLM) with negative binomial distribution. The effect of mass was assessed separately from elytron length and SMI due to collinearity. Models with interaction terms included the corresponding additive terms, but for simplicity the shortcut formula is presented (i.e. a model with y ~ x + z + x:z is presented as y ~ x * z). Model rank, the model structure, model degrees of freedom (*df*), model log-likelihood (*LogLik*), model Aikaike information criterion for small sizes (*AICc*), AICc weight (*ω*) and model coefficient of determination ($R^2$) are shown.

| *Model Rank* | *Model* | *df* | *LogLik* | *AICc* | *ΔAICc* | *ω* | $R^2$ |
|---|---|---|---|---|---|---|---|
| 1 | *Mortality ~ feeding treatment* | 4 | -54.766 | 117.692 | 0.000 | 0.286 | 0.077 |
| 2 | *Mortality ~ feeding treatment + mass* | 5 | -54.619 | 119.480 | 1.788 | 0.117 | 0.078 |
| 3 | *Mortality ~ feeding treatment + elytron length* | 5 | -54.679 | 119.601 | 1.909 | 0.110 | 0.078 |
| 4 | *Mortality ~ feeding treatment + SMI* | 5 | -54.748 | 119.737 | 2.045 | 0.103 | 0.077 |
| 5 | *Mortality ~ feeding treatment + elytron length + SMI* | 6 | -54.607 | 121.554 | 3.862 | 0.041 | 0.079 |
| 6 | *Mortality ~ feeding treatment * elytron length* | 8 | -52.529 | 121.645 | 3.953 | 0.040 | 0.093 |
| 7 | *Mortality ~ feeding treatment * elytron length + feeding treatment*SMI* | 9 | -52.517 | 123.772 | 6.080 | 0.014 | 0.094 |
| 8 | *Mortality ~ feeding treatment * SMI* | 8 | -53.855 | 124.298 | 6.606 | 0.010 | 0.084 |
| 9 | *Mortality ~feeding treatment * mass* | 8 | -53.964 | 124.515 | 6.823 | 0.009 | 0.083 |
| 10 | *Mortality ~ feeding treatment*SMI + elytron length* | 9 | -53.699 | 126.135 | 8.443 | 0.004 | 0.085 |
| 11 | *Mortality ~ feeding treatment * elytron length + feeding treatment * SMI* | 12 | -51.531 | 128.357 | 10.665 | 0.001 | 0.101 |
| 12 | *Null model* | 1 | -64.992 | 132.000 | 14.308 | 0.000 | 0.000 |
| 13 | *Mortality ~ elytron length* | 2 | -64.561 | 133.171 | 15.479 | 0.000 | 0.003 |
| 14 | *Mortality ~ SMI* | 2 | -64.726 | 133.500 | 15.808 | 0.000 | 0.002 |
| 15 | *Mortality ~ mass* | 2 | -64.952 | 133.951 | 16.259 | 0.000 | 0.000 |
| 16 | *Mortality ~ elytron length + SMI* | 3 | -64.480 | 135.056 | 17.364 | 0.000 | 0.004 |
